# Mechanical and Heat Hyperalgesia upon Withdrawal from Chronic Intermittent Ethanol Vapor depends on Sex, Exposure Duration and Blood Alcohol Concentration in Mice

**DOI:** 10.1101/2022.08.19.504595

**Authors:** AJ Brandner, AM Baratta, RS Rathod, C Ferguson, BK Taylor, SP Farris

## Abstract

Approximately half of patients with alcohol use disorder (AUD) report pain and this can be severe during withdrawal. However, many questions remain regarding the importance of sex, blood alcohol concentration (BAC), time course, and pain modality. To examine the impact of sex and BAC on the time course of the development of mechanical and heat hyperalgesia, we characterized a mouse model of Chronic Alcohol Withdrawal Induced Pain (CAWIP) in the presence or absence the alcohol dehydrogenase inhibitor, pyrazole. Male and female C57BL/6J mice underwent chronic intermittent ethanol vapor (CIEV) ± pyrazole exposure for 4 weeks, 4 days/week to induce ethanol dependence. Hind paw sensitivity to the plantar application of mechanical (von Frey filaments) and radiant heat stimuli were measured during weekly observations at 1, 3, 5, 7, 24, and 48 hr after cessation of ethanol exposure. In the presence of pyrazole, males developed mechanical hyperalgesia after the first week of CIEV exposure, peaking at 48 hours after cessation of ethanol. By contrast, females did not develop mechanical hyperalgesia until the fourth week; this also required pyrazole and did not peak until 48 hours. Heat hyperalgesia was consistently observed only in females exposed to ethanol and pyrazole; this developed after the first weekly session and peaked at 1 hour. We conclude that CAWIP develops in a sex –, time –, and BAC – dependent manner in C57BL/6J mice.

## 1 Introduction

Alcohol use disorder (AUD) is characterized as compulsive intake of alcohol, binge drinking to high levels of intoxication (Carvalho et al., 2019; Edward and Koob, 2010), and physiological dependence (Becker, 2008; Koob & Le Moal, 2008). Dependence manifests as an alcohol withdrawal syndrome that includes negative emotional states when access to alcohol becomes limited or cut off completely (Heilig et al., 2010). Key negative emotional states that causally determine the escalation of casual drinking behavior to uncontrolled alcohol consumption are thought to include pain. Pain is a hallmark symptom in 43–73% of individuals with AUD, and manifests as increased hyperalgesia during alcohol withdrawal due to sensitized central and peripheral mechanisms (Apkarian et al., 2013; Edwards et al., 2020; Maleki et al., 2019; Robins et al., 2019; Zale et al., 2015). Conversely, escalation of alcohol consumption contributes to the development of hyperalgesia and chronic pain (Pahng & Edwards, 2021). Individuals with AUD often drink to relieve or prevent alcohol withdrawal-induced pain (Ditre et al., 2019; Egli et al., 2012; Jakubczyk et al., 2016; Maleki et al., 2019; Witkiewitz et al., 2015). Thus, prevention of withdrawal-induced pain may help individuals with AUD remain abstinent from alcohol. However, a better understanding of the mechanisms that lead to the development and maintenance of pain in AUD patients is needed. To address this gap in knowledge, we developed and characterized a mouse model of Chronic Alcohol Withdrawal Induced Pain (CAWIP). We focused on chronic intermittent ethanol vapor (CIEV) paradigms, “the gold standard” for the study of the physical signs of alcohol dependence: (Gilpin et al., 2009; Goldstein, 1972; Vendruscolo & Roberts, 2014).

In rodents, repeated cycles of ethanol exposure (in the diet, by oral gavage, or by vapor) and withdrawal are necessary to model alcohol dependence and the development of persistent negative affective states (Becker & Lopez, 2004; Gilpin et al., 2009; Griffin III, Lopez, Yanke, et al., 2009; Rogers et al., 1979; Vendruscolo & Roberts, 2014). Alcohol withdrawal includes mechanical and heat hyperalgesia (Alongkronrusmee et al., 2016; Avegno et al., 2018; De Logu et al., 2019; Dina et al., 2000; Edwards et al., 2012; Fu et al., 2015; Gatch and Selvig, 2002; Kang et al., 2019; Pradhan et al., 2019; Quadir et al., 2021; Roltsch Hellard et al., 2017; Smith et al., 2017). However, these studies have not systematically considered key factors, including the impact of sex, ethanol exposure duration, time of testing after cessation of ethanol, nor blood alcohol concentration on the intensity of pain-like behaviors.

Previous studies have failed to incorporate females, despite the fact that females generally exhibit greater distress from painful stimuli, greater sensitivity to experimentally induced pain, and a weaker descending control of pain when compared to males (Paller et al., 2009; Popescu et al., 2010). In addition, women have a higher risk of exposure to alcohol than men during adolescence, which translates to higher occurrence and severity of AUD during adulthood (Foster et al., 2015, Foster 2018). Despite these clinically relevant differences in pain sensitivity and AUD between males and females, previous studies have not examined sex – related differences on mechanical and heat sensitivities caused by chronic alcohol exposure and withdrawal. To address this gap, we evaluated CAWIP in both male and female mice.

Also, for the first time, we tracked behavior across multiple weekly sessions of CIEV, evaluated multiple modalities of hypersensitivity (both mechanical and heat), and measured hyperalgesia at several timepoints after the initiation of ethanol withdrawal. Finally, we conducted our studies in the presence or absence of the alcohol dehydrogenase inhibitor, pyrazole, which is well known to dramatically increase blood alcohol levels in mice.

## 2 Methods

### 2.1 Animal Husbandry

64 male and 64 female C57BL/6J mice were obtained from Jackson Laboratories (Bar Harbor, ME) and housed in a temperature (22 – 25°C) and humidity (30 – 70%) controlled room on a 12 – hour light/dark cycle (lights on at 7:00) with *ab libitum* access to food and water. Body weight was recorded twice a week from arrival to euthanasia. Upon arrival, mice acclimated to the facility in their home cage for at least one week. All protocols were approved by the Institutional Animal Care and Use Committee at the University of Pittsburgh and experiments were conducted in accordance with the National Institutes of Health Guidelines for the Care and Use of Laboratory Animals.

### 2.2 Chronic Intermittent Ethanol Exposure (CIEV)

32 mice of each sex underwent exposure to alcohol vapor as previously described (Goldstein, 1972). Each of the four sessions (Monday–Thursday) included 16 hours of exposure to ethanol (17:00 – 9:00), followed by 8 hours of exposure to ambient room air (9:00 – 17:00). 32 mice of each sex served as Air – Control (AC) groups and were moved between housing racks and the induction chambers at 9:00 and 17:00 as were the CIEV groups. The temperature and humidity inside the chambers were maintained at 22 – 25°C and 30 – 70%, respectively. Circulating ethanol vapor levels within the chambers were monitored with a custom voltage sensor generously provided by Brian McCool (Wake Forest University). Cohorts were split into two main groups: alcohol exposed or air – control. The ethanol exposed group was further divided into two groups that received just ethanol (1.5 g/kg in saline, i.p Decon Labs, King of Prussia, PA) or ethanol (1.5 g/kg) with the alcohol dehydrogenase inhibitor pyrazole (68 mg/kg, i.p.; Sigma – Aldrich, P56607 – 6G; **Figure 1A**). The air – control group was also further divided into two separate groups that received either pyrazole (68 mg/kg in saline) or just saline (Gibco, Waltham, MA; **Figure 1A**). After pre – CIEV or air – control treatment, animals were immediately placed into the ethanol induction or control chambers **(Figure 1B)**. Three days of behavioral testing (Friday – Sunday) began the morning after the last CIEV session **(Figure 1C)**. This pattern of 4 days of CIEV followed by 3 days of behavioral testing was continued for 4 weeks **(Figure 1D)**.

**Figure 1.**
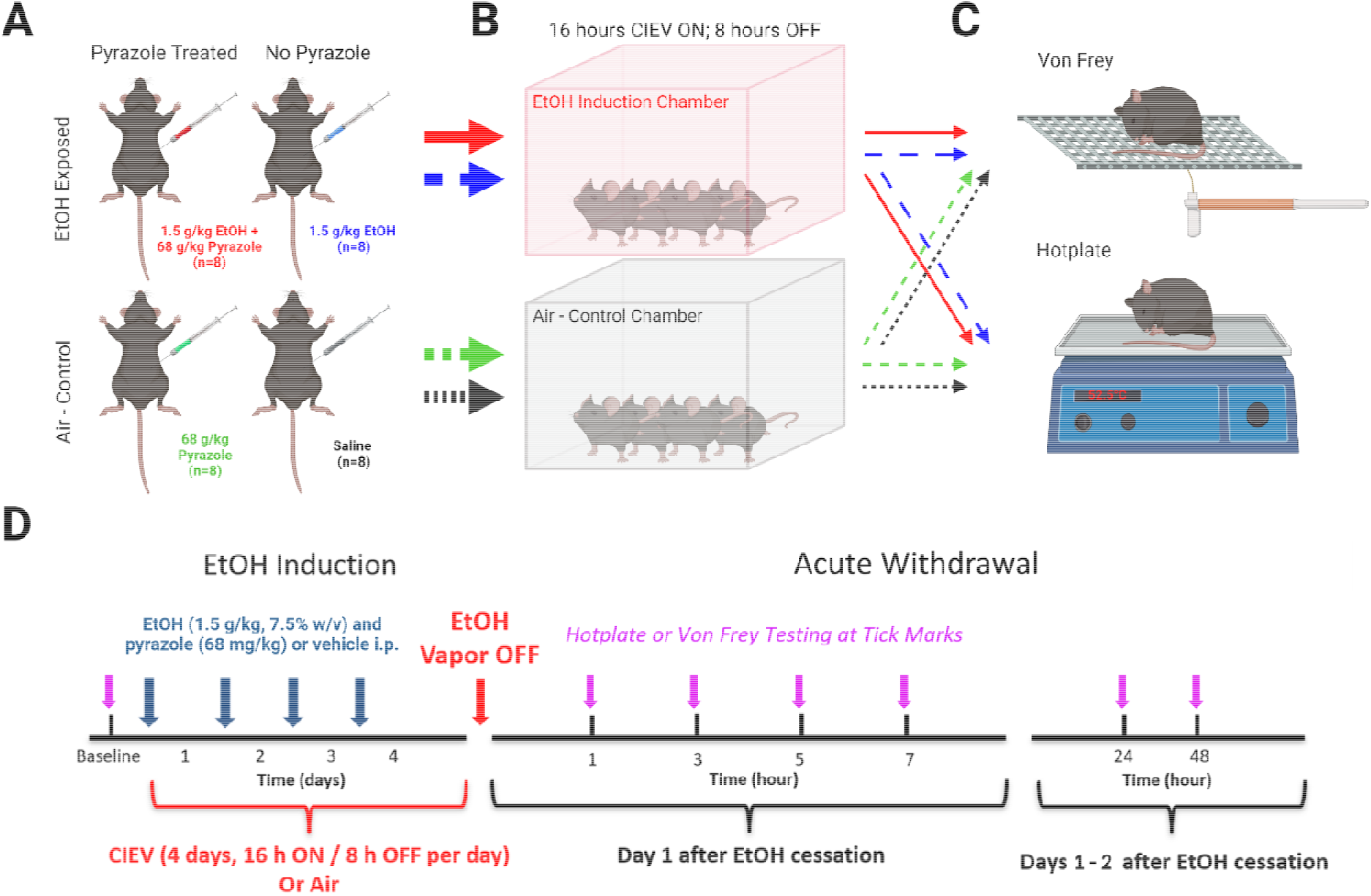
Graphical illustration of methods and timeline used in the current study. **(A)** Mice were separated into four groups and treated with 1.25 mg/kg EtOH + 1 mmol/kg pyrazole, 1.25 mg/kg EtOH, 1 mmol/kg pyrazole, or saline. **(B)** EtOH treated mice were placed into vapor chambers and exposed to 16 hours of EtOH vapor followed by a period of 8 hours of ambient room air. AC mice were also placed into a different vapor chamber without the presence of EtOH. **(C)** Four mice from each treatment group were used for either hotplate or von Frey behavioral testing. **(D) Schematic representation of the EtOH exposure and behavioral timeline**.

Each week, blood alcohol concentrations (BACs) were assessed to ensure our experimental animals were experiencing dependent – like levels of alcohol. Blood was taken via tail vein after the third cycle of CIEV each week. Blood was centrifuged at 2300 x *g* for 5 minutes and plasma was analyzed with an Analox Alcohol Analyzer (AM1, Analox Instruments, London, UK). Ethanol flow rate into the chamber was adjusted to maintain BACs between 175 – 225 mg/dL in the pyrazole – treated CIEV group.

### 2.3 Sensory Testing

Mice were acclimated to either the hotplate or von Frey apparatuses in a temperature and light controlled room for one hour a day between 9:00 – 11:00 for three days, Friday through Sunday. For hotplate acclimation, mice on the first day were placed onto the apparatus at room temperature. On the second and third days, mice were placed onto the apparatus at 52.5ºC and latency to response was recorded. Response latency typically decreased during these sessions, but then stabilized before baseline testing. Baseline testing commenced the following Monday before the first CIEV session. Upon removal from the CIEV chamber, mice were moved to the hotplate or von Frey apparatus and were tested after cessation of alcohol vapor at 1, 3, 5, 7, 24, and 48 hour timepoints, as many withdrawal symptoms in rodents occur 24 – 48 hours after cessation of alcohol (Heilig et al., 2010).

#### Hotplate Test

Mice were placed on a 10” x 10” hotplate heated to 52.5ºC, enclosed within a 10” x 16” acrylic chamber (Columbus Instruments, Columbus, OH). Upon observation of a jump, or rapid flicking or licking a hind paw, the mouse was immediately removed, and response latency recorded (Deuis et al., 2017; Nelson et al., 2019). If no response was made within 30 seconds after placing mouse onto the hotplate, the animal was removed to avoid lasting tissue injury. Three trials were conducted at 10 – minute intervals and averaged.

#### Von Frey Test

Mice were placed on a wire–mesh rack within individual 4” x 4’ x 14” Plexiglas containers. After an acclimation period of at least 30 minutes, a set of eight monofilaments at logarithmic intervals from 0.008 to 6 grams (0.008, 0.023, 0.07, 0.16, 0.4, 1, 2, and 6 grams; Stoelting, Wood Dale, IL) were applied to the center of the plantar surface of the left hind paw for up to 4 seconds using the up–down method (Chaplan et al., 1994). A withdrawal response was defined by rapid retraction of the paw unrelated to normal ambulation. 50% thresholds were calculated using methods as described by Chaplan et. al., and Dixon (Chaplan et al., 1994; Dixon, 1980).

### 2.4 Statistical Analysis

Graphpad Software version 9.3.0 (La Jolla, CA) was used for graphical presentation and statistical analysis. Mechanical thresholds and heat latency data were analyzed separately. Data were first collapsed across Session (Weeks 1 – 4) and Time (hours 1 – 48 hours after cessation of treatment), and then analyzed the main effects of Treatment x Sex with two – way ANOVA; significant main effects were followed by post-hoc multiple comparisons. We then evaluated male and female data separately as well as weekly Session 1 – 4 data separately -- this enabled Treatment x Time two – way repeated measures ANOVA; significant main effects were followed by Sidak’s multiple comparison tests (for data collapsed across Session and Time) or Bonferroni *post-hoc* analysis (for data segregated by Sex and Session). Mann Whitney tests were used to analyze data between two groups at specific timepoints during alcohol withdrawal. Statistical significance was set at p < 0.05. All data are presented as mean ± SEM. Degrees of freedom F and p values for the Treatment x Time two – way repeated measures ANOVA are reported in **Supplemental Table 1**.

## 3 Results

### 3.1 Withdrawal from CIEV ± Pyrazole increases Mechanical and Heat Sensitivity

We first evaluated the data when collapsed (averaged) across Session (Weeks 1 – 4) and Time (hours 1 – 48 after cessation of treatment), to focus on the overall effect of withdrawal from ethanol vapor exposure on mechanical and heat sensitivity.

#### 3.1.1 Administration of pyrazole alone did not change mechanical or heat sensitivity

To determine if administration of pyrazole itself influenced mechanical or heat sensitivity, **Supplemental Figures 1 & 2** compared the air control groups treated with either pyrazole or saline. Two – way ANOVA revealed no differences in either mechanical or heat sensitivity, regardless of sex (P > 0.05). Therefore, the data of these control groups were combined into one group, henceforth referred to as “Air Control”, or “AC”. This AC group was included in subsequent analyses to determine whether withdrawal from CIEV ± pyrazole resulted in sex-dependent changes in mechanical or heat sensitivity (**Figure 2**).

**Figure 2.**
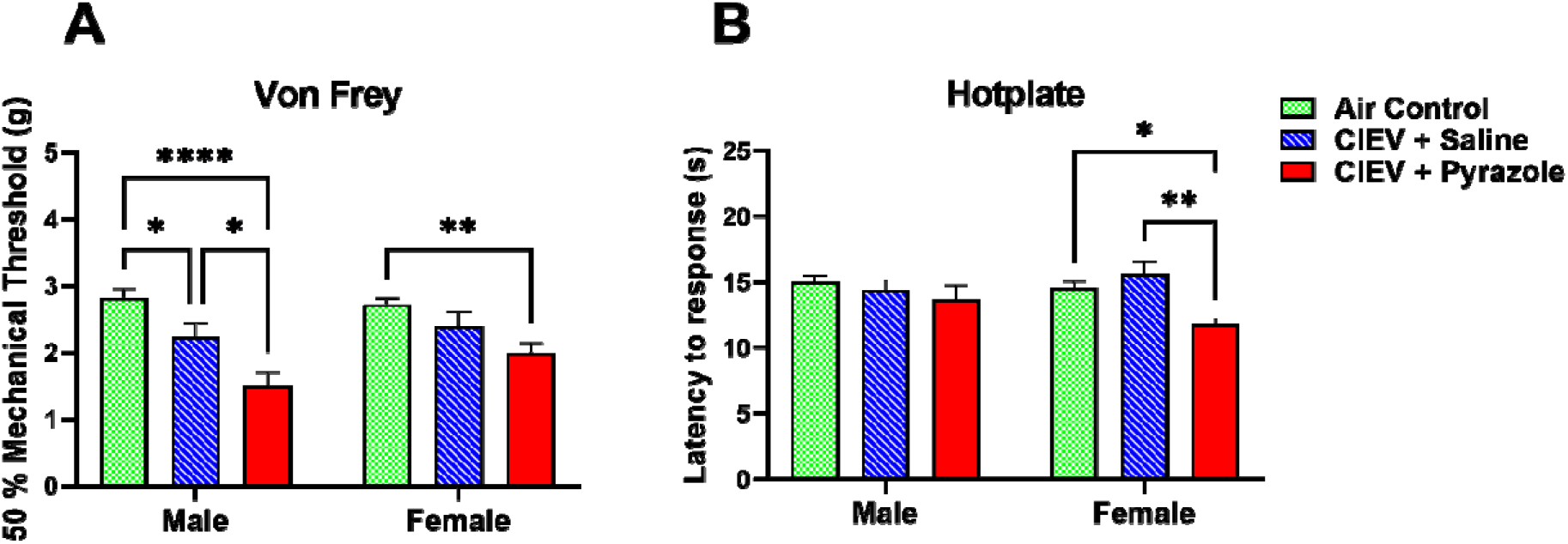
Effect of Chronic Intermittent Ethanol Vapor on von Frey thresholds and hotplate response latencies in males and females. Data were averaged across Session (Weeks 1 – 4 of CIEV) and Time (Hours 1 – 48 after cessation of CIEV). **A)** CIEV + Pyrazole cessation decreased mechanical sensitivity in both sexes. **B)** CIEV + Pyrazole cessation decreased but heat sensitivity in female but not male mice. All data are presented as mean ± SEM. N=8 in each CIEV group; N=16 in Air Control (AC) group. * P < 0.05 CIEV + pyrazole compared to AC, CIEV + Saline compared to AC, or CIEV + Pyrazole compared to CIEV + Saline; ** P < 0.01 CIEV + Pyrazole compared to AC or CIEV + Pyrazole compared to CIEV + Saline; **** P < 0.001 CIEV + pyrazole compared to AC.

#### 3.1.2 Mechanical hypersensitivity in both sexes

Two – way ANOVA (Treatment x Sex) revealed a main effect of Treatment (F (2, 58) = 22.82, P < 0.001). **Figure 2** illustrates that CIEV + Pyrazole cessation decreased mechanical sensitivity in each sex, with reductions in von Frey thresholds of 47% in males and 27% in females when compared to the AC groups, and of 36% in males when compared to the CIEV + Saline group (p < 0.05 by multiple comparisons).

#### 3.1.3. Heat hypersensitivity only in females

In females, two-way ANOVA (Treatment x Sex) revealed a main effect of Treatment (F (2, 58) = 5.84, P < 0.01). CIEV + Pyrazole withdrawal decreased hotplate response latencies by 19% when compared to the AC group, and by 24% when compared to the CIEV + Saline group (p < 0.05 by multiple comparisons). In males, CIEV + Pyrazole did not change hotplate latency (p > 0.05).

### 3.2. Time course of CIEV withdrawal-induced mechanical and heat hypersensitivity

We next determined the time course of mechanical and heat hypersensitivity with a week-by-week analysis of each cycle of CIEV and withdrawal (**Figures 3 – 4**). Statistical results are reported in **Supplemental Table 1**.

**Figure 3.**
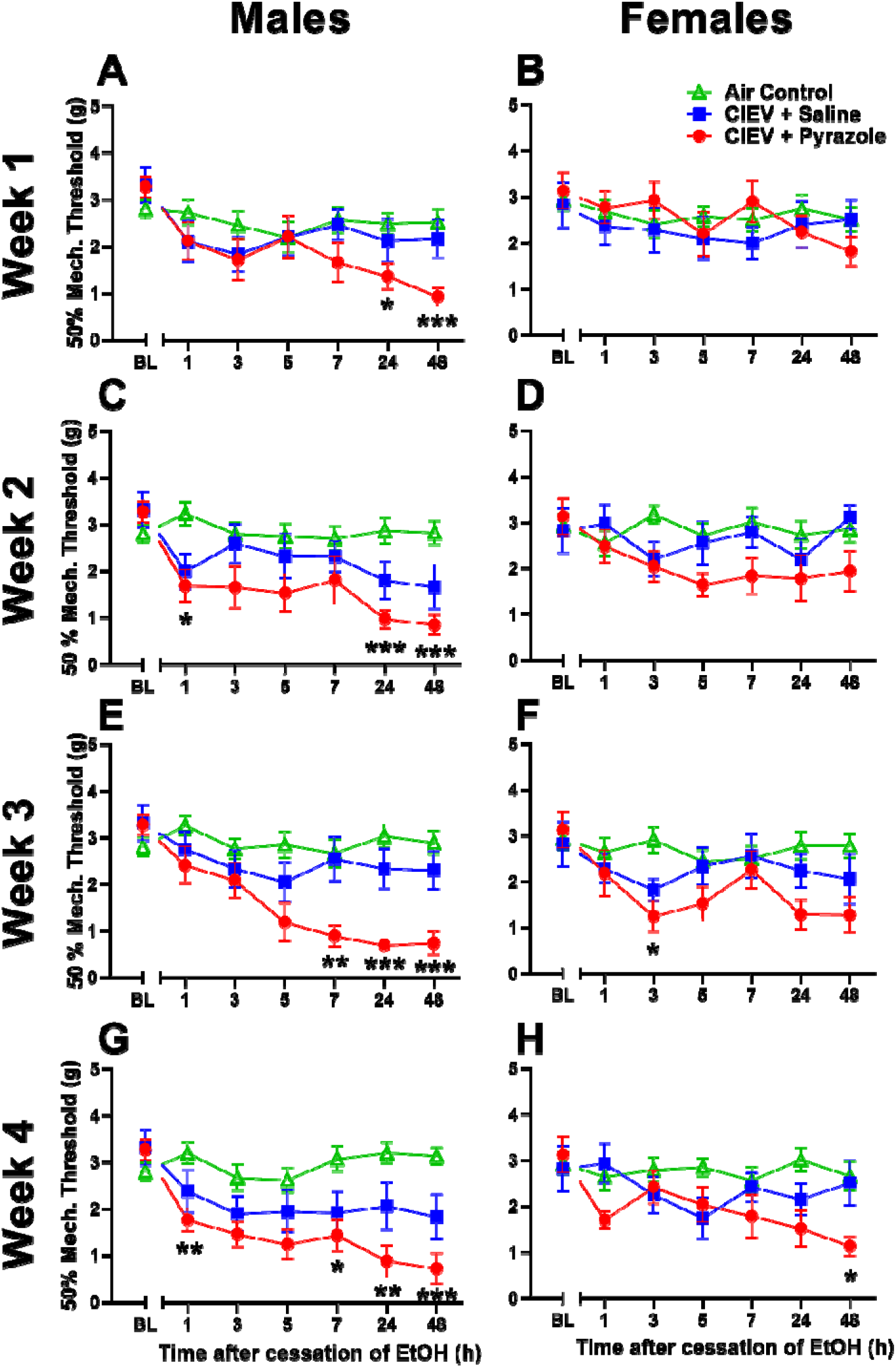
Mechanical thresholds for males (A, C, E, G) and females (B, D, F, H) across the withdrawal period during week 1 (A, B), week 2 (C, D), week 3 (E, F) and week 4 (G, H). Male mechanical sensitivity begins after one week of treatment of CIEV and becomes maximal after four weeks while the females first experience mechanical sensitivity after the third week of CIEV treatment. All data are presented at mean ± SEM. N = 8 per EtOH exposed group, and 16 per AC group. * P < 0.05 CIEV + Pyrazole compared to AC; ** P < 0.01 CIEV + Pyrazole compared to AC; *** P < 0.001 CIEV + Pyrazole compared to AC.

#### 3.2.1 Week 1

##### Mechanical Sensitivity

In males, two-way ANOVA (Treatment x Time) revealed a main effect of Time. CIEV + Pyrazole cessation decreased mechanical threshold compared to the AC group (p<0.05 by multiple comparisons), a decrease of 64% **(Figure 3A)**. Mechanical thresholds gradually decreased in the CIEV + Pyrazole group at hours 24 and 48 when compared to AC, with a peak effect of 63% at the 48 – hour timepoint. In females, no differences were observed (**Figure 3B)**.

##### Heat Sensitivity

We found main effects of Time, Treatment, and a Time x Treatment interaction, indicating that the effect of ethanol withdrawal was time-dependent in females **(Figure 4B)**. CIEV + Pyrazole cessation decreased heat latency by an average of 27% when compared to the AC group at hour 3 and 5, with a peak effect of 29% when compared to the CIEV + saline group at hour 1 (p<0.05 by multiple comparisons). In males, no differences were observed (**Figure 4A)**.

**Figure 4.**
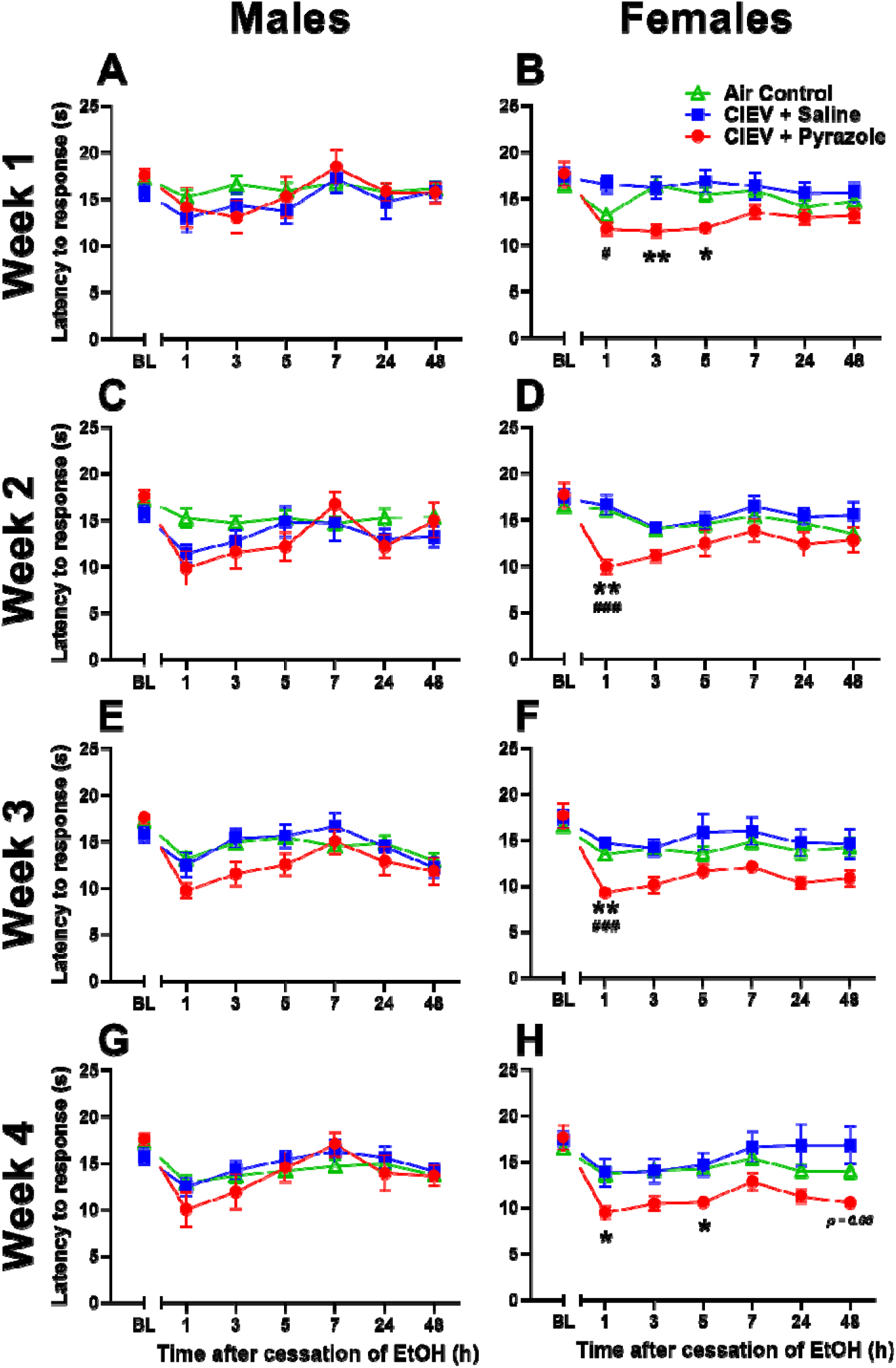
Heat response latencies for males (A, C, E, G) and females (B, D, F, H) across the withdrawal period during week 1 (A, B), week 2 (C, D), week 3 (E, F) and week 4 (G, H). Female heat sensitivity begins after one week of treatment of CIEV and becomes maximal after four weeks while the males do not develop heat sensitivity after CIEV treatment. All data are presented at mean ± SEM. N = 8 per EtOH exposed group, and 16 per AC group. * P < 0.05 CIEV + Pyrazole compared to AC; ** P < 0.01 CIEV + Pyrazole compared to AC.

#### 3.2.2 Week 2

##### Mechanical Sensitivity

In males, two–way ANOVA revealed main effects of Time, Treatment, and a Time x Treatment interaction, indicating that the effect of ethanol withdrawal was time-dependent. CIEV + Pyrazole cessation gradually decreased mechanical threshold compared to the AC group at hours 1, 24, and 48 during the withdrawal period (p<0.05 by multiple comparisons), with a peak effect of 70% at the 48 – hour timepoint **(Figure 3D)**. In females, no differences in mechanical thresholds were detected.

##### Heat Sensitivity

We found a main effect of Treatment and a Treatment x Time interaction in female mice, indicating that the effect of ethanol withdrawal was time-dependent **(Figure 4D)**. CIEV + Pyrazole cessation decreased heat response latency by 39% when compared to the AC group during hour 1 of withdrawal, and by 40% as compared to the CIEV + Saline group during hour 1 of withdrawal (p<0.05 by multiple comparisons). We report a Treatment x Time interaction in males, but multiple comparisons test did not confirm differences at specific time points. To avoid a type II error, further analysis was conducted with non – parametric Mann Whitney tests. These revealed a significant decrease in heat response latency in the male CIEV + Pyrazole group when compared to AC during hour 1 of withdrawal (CIEV + Pyrazole: mean = 9.91 ± 1.74, median = 10.45; AC group: mean = 15.24 ± 1.02, median = 15.15; P < 0.05 by two – tailed Mann Whitney test).

#### 3.2.3 Week 3

##### Mechanical Sensitivity

In males, two-way ANOVA revealed main effects of Treatment, Time, and a Treatment x Time interaction, indicating that the effect of ethanol withdrawal was time dependent. CIEV + Pyrazole cessation rapidly decreased mechanical threshold compared to the AC group at hours 7, 24, and 48 (p<0.05 by multiple comparisons), with a peak effect of 77% at the 24 – hour timepoint **(Figure 3E)**. Females exhibited main effects of Treatment and Time, and CIEV + pyrazole decreased mechanical thresholds by 58% during the 3rd hour of withdrawal **(Figure 3F)**.

##### Heat Sensitivity

We found main effects of Treatment and a Treatment x Time interaction in females. CIEV + Pyrazole cessation decreased heat response latency by 31% as compared to AC and by 37% as compared to CIEV + saline during hour 1 of withdrawal (p < 0.05 by multiple comparisons; **Figure 4F)**. In males, we found a Treatment x Time interaction that could not be confirmed with multiple comparisons test using the Bonferroni correction. Subsequent non – parametric tests revealed a significant decrease in heat response latency in the male CIEV + Pyrazole group when compared to AC during hour 1, 3, and 5 of withdrawal (Hour 1: CIEV + Pyrazole mean = 9.76 ± 0.8, median = 9.60; AC mean = 13.20 ± 0.70, median = 13.71, p < 0.01. Hour 3: CIEV + Pyrazole mean = 11.58 ± 1.30, median = 11.30; AC mean = 14.94 ± 0.59, median = 14.80; p < 0.05. Hour 5: CIEV + Pyrazole: mean = 12.58 ± 1.16, median = 12.15; AC mean = 15.52 ± 0.62, median = 15.70, p < 0.05. All by two – tailed Mann Whitney test).

#### 3.2.4 Week 4

##### Mechanical Sensitivity

In males, two-way ANOVA revealed a main effect of Treatment, Time, and a Treatment x Time interaction, indicating that the effect of alcohol withdrawal was time dependent. CIEV + Pyrazole cessation rapidly decreased mechanical threshold compared to the AC group at hours 1, 7, 24, and 48 (p<0.05 by multiple comparisons), with a peak effect of 77% at the 48 – hour timepoint **(Figure 3G)**. In females, we observed main effects of Treatment, Time, and a Treatment x Time interaction, and CIEV + pyrazole gradually decreased mechanical thresholds by 57% when compared to AC at hour 48 during withdrawal **(Figure 3H)**.

##### Heat Sensitivity

We found main effects of Treatment, Time, and a Treatment x Time interaction in females. CIEV + Pyrazole cessation decreased heat response latency as compared to the AC control group in hours 1 and 5 during withdrawal, with a peak effect of 30% at the 1 – hour timepoint. **(Figure 4H)**. There were no main effect of CIEV + pyrazole in males.

### 3.3 Blood Alcohol Concentrations

As expected, inhibition of alcohol dehydrogenase with pyrazole dramatically increased BACs to the 150–250 mg/dL range at the end of 16h of exposure in both male and female mice exposed to CIEV **(Supplemental Figure 3**).

## Discussion

Chronic alcohol withdrawal-induced pain (CAWIP) has been studied with a variety of paradigms in rats and mice. We chose a model of chronic intermittent ethanol vapor (CIEV) because it is now recognized as the gold standard in the study of alcohol dependence, allows for precise control of blood alcohol concentration (BAC), and provides reliable and repeatable outcome measures, including mechanical and heat hypersensitivity. Our study establishes the effects of multiple cycles of chronic alcohol vapor exposure and subsequent withdrawal periods on mechanical and heat sensitivity in male and female mice and reveals several important principles in the use of CIEV to study CAWIP. First, we noted that previous studies of alcohol dependence (i.e. alcohol consumption and anxiety-like behaviors upon cessation of alcohol) were restricted to CIEV in the presence of pyrazole (Becker et al., 1997; Griffin III, Lopez, & Becker, 2009; Kliethermes et al., 2004; Littleton et al., 1974). To determine whether pyrazole was necessary for other signs of alcohol dependence (in this case, pain-like behaviors upon cessation of alcohol), we included groups that did not receive pyrazole. We found that pyrazole increased the severity of mechanical hyperalgesia in a “dose response-like” manner and was necessary for the detection of heat hyperalgesia. These results confirm that pyrazole is a necessary component of CIEV protocols to study CAWIP. Second, previous studies were restricted to male rats or mice. We studied both male and female mice, leading to the discovery of two dramatic sex differences: males exhibited greater mechanical hyperalgesia, while only females exhibited heat hyperalgesia. Third, previous studies were restricted to analysis of just one session of pain-like behaviors after cessation of CIEV. To determine the developmental time course of CAWIP, we evaluated behavior on four consecutive weeks. We found that mechanical hyperalgesia peaked after 3 weekly cycles of CIEV, and this continued for at least one additional week.

### Pyrazole is necessary for the full manifestation of CAWIP

Pyrazole is an alcohol dehydrogenase inhibitor that inhibits the metabolism of alcohol into acetaldehyde. In mouse models of CIEV, pyrazole is required to achieve BACs at levels necessary to achieve dependence (Becker & Lopez, 2004; Griffin III, Lopez, Yanke, et al., 2009). Pyrazole alone can produce toxic effects in mice such as weight loss and liver necrosis when combined with alcohol (Goldstein & Pal, 1971; Lelbach, 1969). One group found a strong withdrawal phenotype in CIEV treated mice without the usage of pyrazole, suggesting that pyrazole is not needed in mouse CIEV studies (Eisenhardt et al., 2015). We found that pyrazole substantially increased BACs in the setting of CIEV, while treatment of the AC + Pyrazole group showed no differences in body weight and pain sensitivity when compared to the AC + Saline group in both males and females. Importantly, pyrazole increased the severity of mechanical hyperalgesia in a “dose response-like” manner and was necessary for the detection of heat hyperalgesia. These results confirm that pyrazole is a necessary component of CIEV protocols to study CAWIP. Our CIEV + Saline groups achieved significantly lower BALs throughout the experiment when compared to the CIEV + Pyrazole groups (Supplemental Figure 3).

### CIEV produced heat hyperalgesia only in female mice

We found that CIEV decreased heat response latency in female but not male mice. By contrast, one previous study (restricted to males only) indicates that chronic alcohol exposure via intermittent access to two – bottle choice (IA2BC) produces heat hypersensitivity 24 hours into the withdrawal period (Quadir et al., 2021). This study used radiant heat to stimulate a paw withdrawal response; by contrast, we placed mice on a hotplate and recorded a response to be represented not only by paw lifting, but also licking or jumping. We do not believe that our contrasting data is a result of different testing modalities since both hotplate and Hargreaves both require the integration of supraspinal pain pathways. (Deuis et al., 2017). Heat hypersensitivity is seen during withdrawal in male rats subjected to liquid diet, IA2BC, and CIEV models of alcohol dependence (Avegno et al., 2018; Dina et al., 2000; Fu et al., 2015; Kang et al., 2019, 2019; Roltsch Hellard et al., 2017). Little data is available showing alcohol withdrawal – induced heat hypersensitivity in male and female mice, however, one study showed no effect of repeated cycles of CIEV and withdrawal on heat sensitivity in both male and female HS/Npt mice (Metten et al., 2018). We conclude that only female mice experience heat sensitivity as a result of CIEV and withdrawal. Sex – dependent mechanisms in heat nociception could explain why we see heat sensitivity in females and not males in our study.

### Males are More Sensitive to Mechanical Stimuli After CIEV

We discovered that male mice exhibit greater mechanical hyperalgesia than female mice. Our results are consistent with previous reports of mechanical sensitivity after withdrawal from chronic alcohol exposure, albeit restricted to just male mice. (Alongkronrusmee et al., 2016; De Logu et al., 2019; Quadir et al., 2021; Smith et al., 2017). Male C57BL/6 mice experienced mechanical sensitivity 24 and 48 hours into withdrawal from alcohol administered via oral gavage, and this sensitivity become more pronounced after each subsequent week of alcohol administration (Alongkronrusmee et al., 2016). This result is similar to those from our study where multiple cycles of CIEV, and withdrawal produced greater mechanical sensitivity. In a IA2BC paradigm, mechanical sensitivity was seen 24 hours into the withdrawal period in male C57BL/6J mice (Quadir et al., 2021; Smith et al., 2017). We can confirm mechanical sensitivity at the 24 – and 48 – hour withdrawal period in the CIEV model of alcohol dependence. Little data is available showing mechanical sensitivity in female mice using any model of alcohol dependence, and we are the first to directly compare mechanical thresholds in male and female mice after repeated cycles of CIEV and withdrawal. In a rat model of CAWIP, one study showed decreased mechanical thresholds in female rats 1 week after discontinuation of a nine weeks of alcohol administration via liquid diet (Cucinello-Ragland et al., 2021). However, they did not test for mechanical sensitivity during the critical 24 – and 48 – hour withdrawal period where physiological signs of dependence are maximal. Overall, multiple cycles of CIEV and withdrawal produced robust mechanical sensitivity in our male mice during the 24 – and 48 – hour of the withdrawal period, while females experienced mechanical sensitivity during the 48 – hour of the withdrawal period.

### Development of Mechanical and Heat Sensitivity during repeated sessions of CIEV

Previous studies were restricted to analysis of just one session of pain-like behaviors after cessation of CIEV. To determine the developmental time course of CAWIP, we evaluated behavior on four consecutive weeks, using a repeated-measures design that that evaluated multiple time points within a 48–hour withdrawal period. We found that male mice rapidly developed mechanical hypersensitivity after one cycle of CIEV, whereas female mice did not exhibit mechanical hypersensitivity until the latter weeks. Mechanical sensitivity in male mice increased after each subsequent week of CIEV and peaked after three weeks. This pattern reflects what is seen in voluntary drinking studies using CIEV in male mice, where voluntary alcohol drinking increases after each subsequent cycle of CIEV and withdrawal. (Griffin III, Lopez, Yanke, et al., 2009; Lopez & Becker, 2005). However, it took the females three cycles of CIEV to experience mechanical sensitivity initially and did not increase after the subsequent fourth week. We conclude that mechanical hyperalgesia peaked after 3 weekly cycles of CIEV, and this continued for at least one additional week in both sexes.

Overall, heat sensitivity is seen in female mice, but not males, after chronic alcohol exposure via treatment with CIEV and Pyrazole. Female mice experienced heat sensitivity during early withdrawal timepoints across all four weeks. The male counterparts did not due to high variability in their behavior. During week 2 and 3, heat sensitivity in the male group is present but was not picked up using conservative statistical methods. Upon further analysis, non – parametric Mann Whitney tests revealed statistical support for potential heat sensitivity in the male CIEV + pyrazole group when compared to AC males during early withdrawal timepoints. Further analysis needs to be conducted with larger sample sizes to account for variability we saw in our male hotplate data. We believe that three to four cycles of CIEV and withdrawal is enough to produce a dependent–like phenotype in male and female C57BL/6J mice, as seen through increased mechanical and heat sensitivity.

## Conclusion

This is the first report of sex differences in CIEV withdrawal – induced pain. We conclude that male and female mice experience dramatic differences in mechanical and heat hypersensitivity. Future studies will identify key neurochemical systems and brain regions that can contribute to CAWIP and to target these with pharmacological intervention. In summary, we demonstrate that four weeks of CIEV is sufficient to produce mechanical hyperalgesia in both male and female C57BL/6J mice, and heat hyperalgesia in female C57BL/6J mice. The pain associated with chronic alcohol use disorder is incredibly debilitating and disrupts cognitive function, and so it will be imperative to test the affective component of CAWIP, as well as to study the different supraspinal mechanisms by which this occurs.

## Acknowledgements

We gratefully acknowledge the support of NIAAA, NIDA, and NINDS grants AA024836 (SPF), AA020889 (SPF), AA029942 (AMB), DA037621 (BKT), NS045954 (BKT), NS112632 (BKT), NS073548 (AJB).

