## Supplemental Table 1 for "Mechanical and Heat Hyperalgesia upon Withdrawal from Chronic Intermittent Ethanol Vapor depends on Sex, Exposure Duration and Blood Alcohol Concentration in Mice"

|  | | | Week 1 | Week 2 | Week 3 | Week 4 |
| --- | --- | --- | --- | --- | --- | --- |
| Von Frey | Males | Interaction | F (12, 174) = 1.61  P = 0.0916 | F (12, 174) = 3.05  **P = 0.0006** | F (12, 174) = 4.55  **P < 0.0001** | F (12, 174) = 4.14  **P < 0.0001** |
|  |  | Time | F (4.383, 127.1) = 5.34  **P < 0.0001** | F (4.366, 126.6) = 7.38 **P < 0.0001** | F (5.199, 150.8) = 9.65  **P < 0.0001** | F (5.268, 152.8) = 8.35  **P < 0.0001** |
|  |  | Treatment | F (2, 29) = 3.24  P = 0.0539 | F (2, 29) = 8.74  **P = 0.0011** | F (2, 29) = 9.68  **P = 0.0006** | F (2, 29) = 12.25  **P = 0.0001** |
|  | Females | Interaction | F (12, 174) = 0.75  P = 0.7032 | F (12, 174) = 1.3  P = 0.2251 | F (12, 174) = 1.52  P = 0.1222 | F (12, 174) = 1.84  **P = 0.0448** |
|  |  | Time | F (5.132, 148.8) = 1.44  P = 0.2131 | F (4.947, 143.5) = 1.66 P = 0.1479 | F (4.716, 136.8) = 3.23 **P = 0.0099** | F (4.840, 140.3) = 2.56  **P = 0.0313** |
|  |  | Treatment | F (2, 29) = 0.59  P = 0.5623 | F (2, 29) = 4.33  **P = 0.0227** | F (2, 29) = 5.65  **P = 0.0085** | F (2, 29) = 4.67  **P = 0.0175** |
| Hotplate | Males | Interaction | F (12, 174) = 1.00  P = 0.4471 | F (12, 174) = 2.52  **P = 0.0044** | F (12, 174) = 2.31  **P = 0.0092** | F (12, 174) = 1.71  P = 0.0687 |
|  |  | Time | F (4.961, 143.9) = 4.91  **P < 0.0004** | F (4.196, 121.7) =8.97 **P < 0.0001** | F (4.470, 129.6) = 16.62 **P < 0.0001** | F (4.471, 129.7) = 13.55  **P < 0.0001** |
|  |  | Treatment | F (2, 29) = 0.60  P = 0.5540 | F (2, 29) = 1.97  P = 0.1577 | F (2, 29) = 1.91  P = 0.1661 | F (2, 29) = 0.2  P = 0.8234 |
|  | Females | Interaction | F (12, 174) = 2.58  **P = 0.0036** | F (12, 174) = 2.75  **P = 0.0019** | F (12, 174) = 1.92  **P = 0.0345** | F (12, 174) = 2.39  **P = 0.0070** |
|  |  | Time | F (3.986, 115.6) = 6.28  **P < 0.0001** | F (4.248, 123.2) = 8.91  **P < 0.0001** | F (4.501, 130.5) = 12.66  **P < 0.0001** | F (3.662, 106.2) = 13.17  **P < 0.0001** |
|  |  | Treatment | F (2, 29) = 4.70  **P = 0.0170** | F (2, 29) = 5.60  **P = 0.0088** | F (2, 29) = 5.19  **P = 0.0119** | F (2, 29) = 5.32  **P = 0.0108** |

**Supplemental Table 1.** Main effects of Treatment (cessation of CIEV ± Pyrazole) and Time (hours after cessation) on responses to mechanical (von Frey) and heat (hotplate) stimuli for each of the 4 weeks of testing. Significant results (p<0.05) are in bold.
