## Supplemental Figure 2 for "Mechanical and Heat Hyperalgesia upon Withdrawal from Chronic Intermittent Ethanol Vapor depends on Sex, Exposure Duration and Blood Alcohol Concentration in Mice"

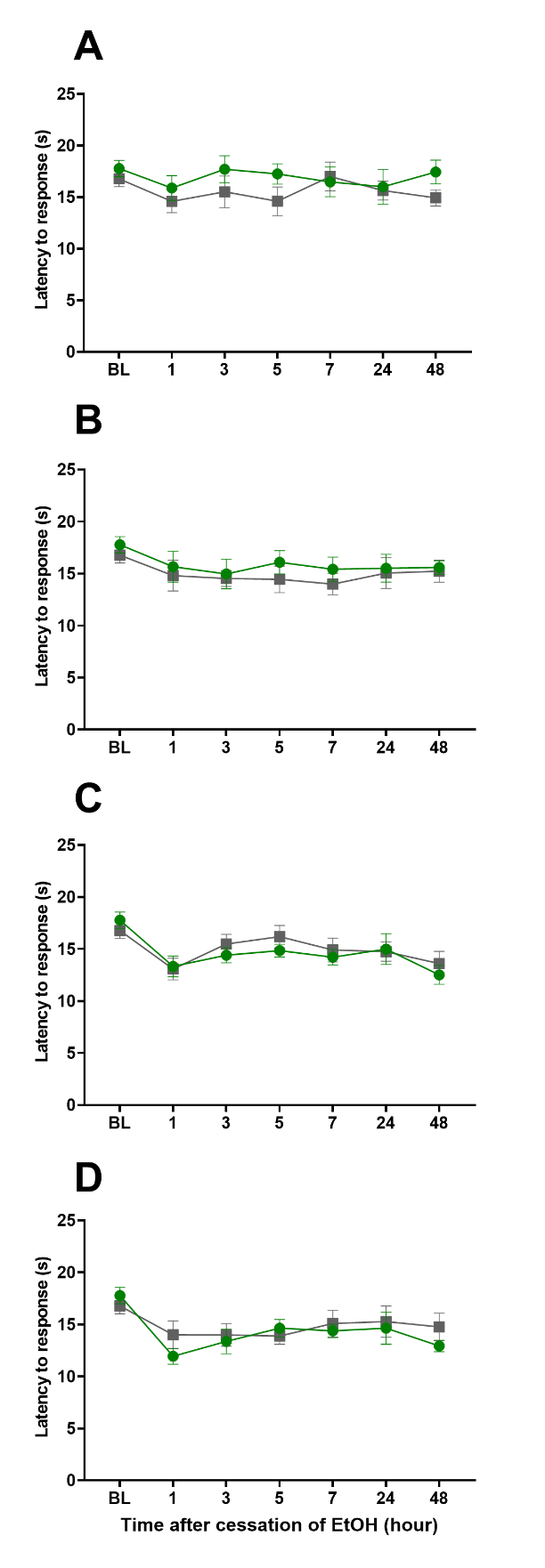

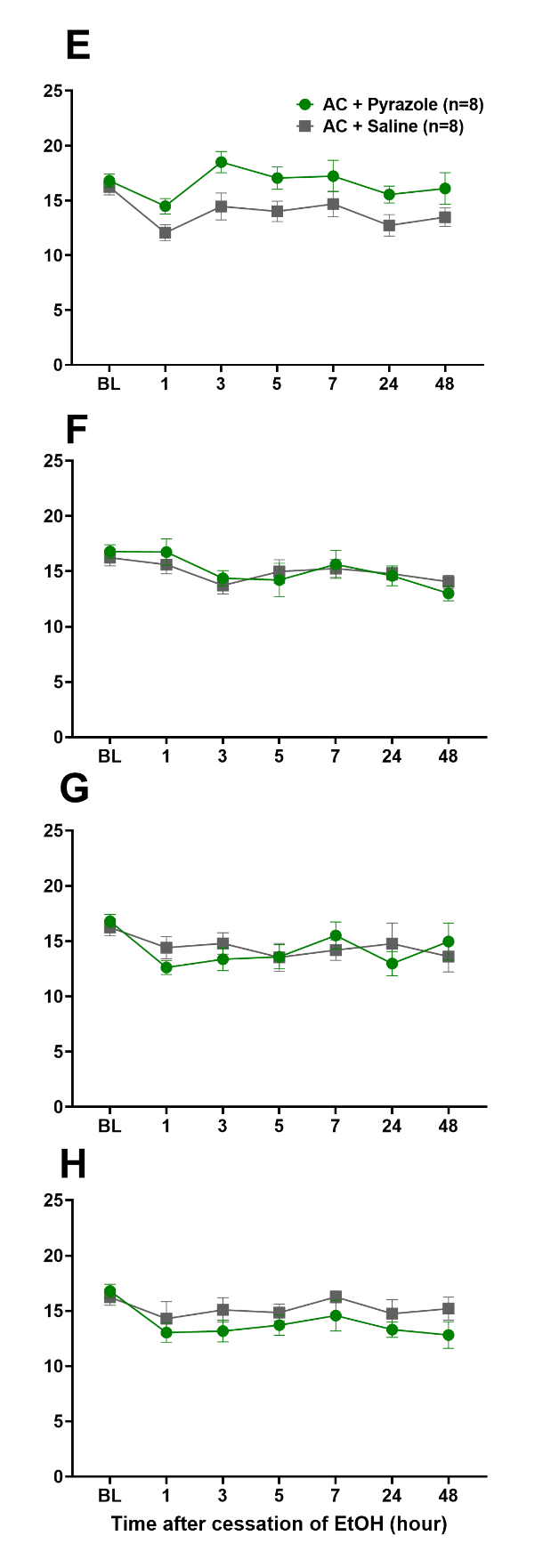

**Supplemental Figure 2** Cessation of pyrazole did not change heat sensitivity. Data for males **(A-D)** and females **(E-H)** for weeks 1 **(A, E)**, 2 **(B, F)**, 3 **(C, G)**, and 4 **(D, H)** of CIEV. All data are presented as mean ± SEM. N = 8 per group.
