## Supplemental Figure 3 for "Mechanical and Heat Hyperalgesia upon Withdrawal from Chronic Intermittent Ethanol Vapor depends on Sex, Exposure Duration and Blood Alcohol Concentration in Mice"

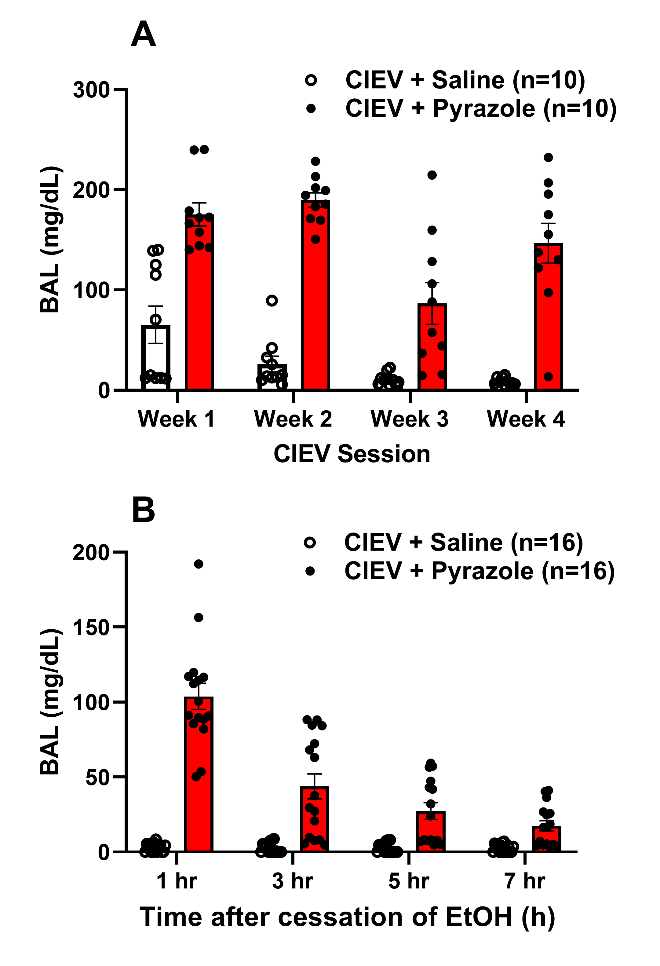

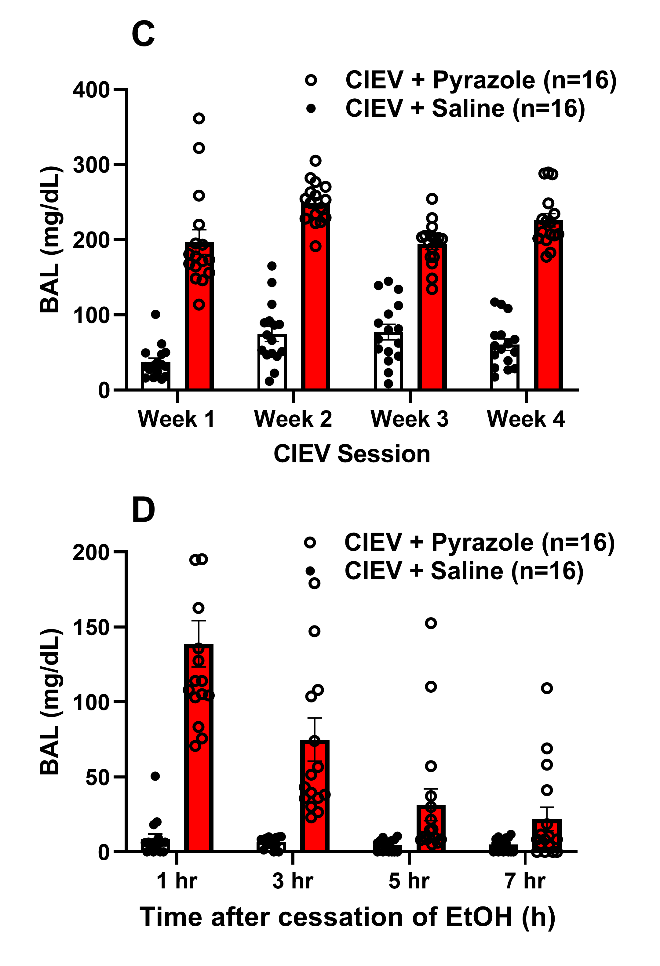

**Supplemental Figure 3.** BACs from male **(A, B)** and female **(C, D)** taken immediately following the last CIEV cycle during weeks 1–4 **(A, C)** and during hours 1, 3, 5, 7 acute withdrawal after week 5 of CIEV **(B, D).** All data are presented as mean ± SEM.
